# Increased drought resistance in state transition mutants is linked to modified plastoquinone pool redox state

**DOI:** 10.1101/2023.03.17.533090

**Authors:** Lucas Leverne, Thomas Roach, François Perreau, Fabienne Maignan, Anja Krieger-Liszkay

**Affiliations:** Université Paris-Saclay, Institute for Integrative Biology of the Cell (I2BC), CEA, CNRS, 91198 Gif-sur-Yvette cedex, France; University of Innsbruck, Department of Botany, Innsbruck, 6020, Austria; Université Paris-Saclay, INRAE, AgroParisTech, Institut Jean-Pierre Bourgin (IJPB), 78000, Versailles, France; Laboratoire des Sciences du Climat et de l’Environnement, LSCE/IPSL, CEA-CNRS-UVSQ, Université Paris-Saclay, Gif-sur-Yvette, France

**Keywords:** photosynthesis, state transitions, plastoquinone redox state, root growth, singlet oxygen

## Abstract

Identifying traits that exhibit improved drought resistance is highly important to cope with the challenges of predicted climate change. We investigated the response of state transition mutants to drought. Compared with the wild type, state transition mutants were less affected by drought. Photosynthetic parameters in leaves probed by chlorophyll fluorescence confirmed that mutants possess a more reduced plastoquinone (PQ) pool, as expected due to the absence of state transitions. Seedlings of the mutants showed an enhanced growth of the primary root and more lateral root formation. The photosystem II inhibitor 3-(3,4-dichlorophenyl)-1,1-dimethylurea, leading to an oxidized PQ pool, inhibited primary root growth in wild type and mutants, while the cytochrome *b*6*f* complex inhibitor 2,5-dibromo-3-methyl-6-isopropylbenzoquinone, leading to a reduced PQ pool, stimulated root growth. A more reduced state of the PQ pool was associated with a slight but significant increase in singlet oxygen production. Singlet oxygen may trigger a, yet unknown, signaling cascade promoting root growth. We propose that photosynthetic mutants with a deregulated ratio of photosystem II to photosystem I activity can provide a novel path for improving crop drought resistance.

**Summary Statement:** *Arabidopsis thaliana* mutants affected in state transitions are more drought resistant.

## Introduction

Drought-resistant plants are required to cope with the increase in world population and the challenges of predicted climate change. New traits need to be identified and methods have to be developed to maintain and improve crop productivity under harsh environmental conditions. Water and nutrients are absorbed by plant roots and specific root traits such as growth and architecture are important under unfavorable conditions, such as drought. Previously it has been shown that inhibition of the carotenoid biosynthesis pathway reduces root growth and the emergence of lateral roots in *Arabidopsis thaliana*, demonstrating the importance of carotenoids for root growth (van Norman et al., 2014). It has been shown that strigolactones and abscisic acid, both being phytohormones derived from carotenoids, regulate root growth and branching of lateral roots (Gomez–Roldan et al., 2008; Ruyter-Spira et al., 2011). Furthermore, the volatile β-cyclocitral that is either enzymatically produced by carotenoid cleavage dioxygenases or by the non-enzymatic ^1^O_2_– dependent oxidation of β-carotene has been found to be a root growth regulator (Dickinson et al., 2019). Beside β-cyclocitral, other carotenoid cleavage products (apocarotenoids) and reactive electrophile species (RES) may be capable of promoting root growth and development (Biwas et al., 2019).

Apocarotenoids are generated either by non-enzymatic or enzymatic carotenoid oxidation. Singlet oxygen (^1^O_2_) oxidizes β-carotene to, amongst others, β-cyclocitral or β-ionone, both substances that are known to be involved in plant growth and development. β-cyclocitral has been shown to induce drought resistance in *Arabidopsis thaliana* (D’Alessandro et al., 2019). In photosystem II (PSII) chlorophyll in its triplet state (^3^Chl) reacts with oxygen, a triplet in its ground state, producing ^1^O_2_. ^3^Chl is generated by charge recombination reactions of the primary radical pair in PSII (Rutherford and Krieger-Liszkay, 2001; Krieger-Liszkay, 2005). Charge recombination leads to the formation of the primary radical pair P^+^Ph^−^, with pheophytin (Ph) being the primary electron acceptor in PSII, and subsequently to the repopulation of the excited state of P680. The excited state formed can be either a singlet or a triplet state (^3^P680), depending on the charge recombination route. The probability of charge recombination within PSII increases when the plastoquinone (PQ) pool and the acceptor side of PSII are reduced, for example, when there is a lack of stromal electron acceptors under CO_2_ limitation from drought-induced stomatal closure.

The redox state of the PQ pool is also influenced by movements of the light-harvesting antenna trimer (L-LHCII), which changes the absorption cross-sections of PSII and PSI in a process called state transition. Both phosphorylation (Depège et al., 2003; Bellafiore et al., 2005) and N-terminal acetylation (Koskela et al., 2018) are posttranslational modifications of proteins involved in this process. Phosphorylation and acetylation of L-LHCII are required to allow the detachment of L-LHCII from PSII to adjust the amount of excitation energy received by the two photosystems (Koskela et al., 2018). These posttranslational modifications are highly dynamic and are often regulated by changes in the environment and stress (e.g., Stone and Walker, 1995; Linster and Wirtz, 2018). When more light is absorbed by PSII than by PSI, L-LHCII attached to PSII becomes phosphorylated and acetylated. The modified L-LHCII detaches from PSII and migrates to PSI, thereby increasing the cross-section and excitation of PSI. Lysine acetylation of L-LHCII is achieved by the chloroplast acetyltransferase enzyme *NSI* (NUCLEAR SHUTTLE INTERACTING; At1G32070) (Koskela et al., 2018). The reduction of the PQ pool regulates the activity of the serine/threonine-protein kinase STN7 (Depège et al., 2003; Bellafiore et al., 2005), while an oxidized PQ pool leads to dephosphorylation of LHCII by the phosphatase PPH1/TAP38 (Pribil et al, 2010; Shapiguzov et al., 2010). N-terminal acetylation has been shown to decrease significantly after drought stress, and transgenic downregulation of this activity induced drought tolerance (Linster et al., 2015). How state transitions are linked to drought tolerance remains unclear.

We hypothesize that a higher reduction state of the PQ pool and concomitant ^1^O_2_ generation leads to a signaling event that induces drought stress tolerance. To test this, we investigated drought resistance of *Arabidopsis thaliana* state transition mutants using the well-characterized mutant *stn7* and the less well-characterized mutants *nsi1* and *nsi2*, two knockout lines lacking the chloroplast acetyltransferase *NSI*.

## Materials and Methods

### Plant material and plant growth conditions

*Arabidopsis thaliana* (ecotype Columbia-0) wild type (WT) *stn7, nsi1*, and *nsi2* were grown in Jiffy-7^®^ – Peat Pellets in plastic pots (5 cm high, 5.5 cm diameter) in ambient air for 4 weeks in a growth cabinet in long-day conditions: 16 h of light (22°C), 8 h of dark (18°C), and at a light intensity of 110 μmol quanta m^−2^s^−1^ (light source, fluorescent light tubes; OSRAM fluora 58 Watt/77). To achieve mild drought stress, watering was stopped for 4-6 days. For each replicate, plants in control and stress conditions of the same age were used.

### Seedling growth conditions

WT, *stn7, nsi1*, and *nsi2* seeds were sterilized in a solution of ethanol 70% (v/v) and sodium dodecyl sulfate 0.05% (w/v) under gentle agitation for 12 min, washed three times with ethanol 96%, and then dried on a sterile filter paper. Seeds were grown in one-half-strength Gamborg’s B5 agar medium (Gamborg et al., 1968). Seeds were stratified in the dark at 4°C for 48 h. Then plates were placed in a growth cabinet 16/8 h day/night, at 22°C/18°C, fluorescent light 110 μmol quanta m^−2^s^−1^. Supplements including 3-(3,4-dichlorophenyl)-1,1-dimethylurea (DCMU, 2 μM), 2,5-dibromo-3-methyl-6-isopropylbenzoquinone (DBMIB, 30 μM), and mannitol (200 mM) were added directly to the agar medium. The stability of DBMIB in the light was tested by measuring the inhibitory effect of the DBMIB solution after 10 days of storage in the growth cabinet under the same photoperiod and temperature as above. Photosynthetic electron transport activity of thylakoids was still inhibited by 80% in the presence of 3 μM DBMIB.

### Relative water content (RWC)

Total rosettes were sampled and weighed. Saturated weight was determined by weighing the rosettes after a 12 h immersion in water at 4°C in the dark. Dry weight was determined by weighing shoots after drying for 48 h at 70°C. RWC was calculated using: RWC = (FW – DW)/(SW – DW) where FW, SW, and DW are the leaf fresh, water-saturated, and dry weights, respectively.

### Stomata opening

In order to observe the stomata under the microscope, transparent tape was applied to the abaxial part of the leaf, and it was quickly torn off so that a thin layer of the lower epidermis could be removed and fixed to be observed under the microscope. The ratio of width/length was determined with an optical microscope × 40 (Zeiss Axio Imager M1), measured manually using the Fiji software, at least for 100 stomata for each condition and each genotype.

### Chlorophyll fluorescence

Chlorophyll fluorescence was measured on whole plants at room temperature with an Imaging-PAM (Walz, Effeltrich, Germany). Plants were adapted for 15 min in the dark before measuring the minimum fluorescence level Fo. A saturating flash (300 ms, 10 000 μmol quanta m^−2^s^−1^) was given to determine the maximum fluorescence Fm. To measure the maximum fluorescence level in the light, Fm’, plants were illuminated for 3 min with different intensities of actinic light (55, 80, 110, 145, 185,230, 335, 425, 610 μmol quanta m^-2^s^-1^), and a saturating flash was given at the end of each light intensity step.

Parameters are defined as follows: F: fluorescence yield; F_0_’: dark fluorescence level after illumination; Fm’: maximal fluorescence yield in the light; NPQ: nonphotochemical fluorescence quenching; Y(II): effective PS II quantum yield; Y(NPQ): quantum yield of regulated energy dissipation in PS II; Y((NO): quantum yield of nonregulated energy dissipation in PS II; qP: photochemical quenching; qL: fraction of open PS II reaction centers. The different fluorescence parameters are calculated as follows: Y(II) = (Fm’-F)/Fm’; NPQ = (Fm-Fm’)/Fm’; Y(NPQ) = 1 -Y(II) - 1/(NPQ+1+qL(Fm/Fo-1)); Y(NO) = 1/(NPQ+1+qL(Fm/Fo-1)); qP=(Fm’-F)/(Fm’-Fo’); qL = qP x Fo’/F.

### 77K Chlorophyll fluorescence measurements

Fluorescence spectra of purified thylakoid were measured with a Carry Eclipse fluorimeter; excitation wavelength: 430 nm. The intensity was normalized to the intensity of the PSI emission.

### Spin-trapping electron paramagnetic resonance (EPR) spectroscopy

^1^O_2_ was trapped using the water-soluble spin probe 2,2,6,6-tetramethyl-4-piperidone hydrochloride (TEMPD-HCl) (Hideg et al., 2011). Thylakoids from Arabidopsis were prepared using a standard protocol (e.g.; Krieger-Liszkay et al., 2019). Thylakoids (20 μg chlorophyll ml^−1^) were illuminated for 2 min with red light (Schott filter RG 630) at 670 μmol quanta m^−2^ s^−1^ in 100 mM TEMPD-HCl 0.3 M sorbitol, 50 mM KCl, 1 mM MgCl2 and 25 mM HEPES (pH 7.6). Spin-trapping assays for detecting ^•^OH derived from O ^• –^/H_2_ O_2_ were carried out using the spin trap 4-pyridyl1-oxide-N-tert-butylnitrone (4-POBN). Leaves were vacuum-infiltrated with a buffer (25 mM HEPES, pH 7.5, 5 mM MgCl_2_, 0.3 M sorbitol) containing the spin trap reagents (50 mM 4-POBN, 4% ethanol, 50 μM Fe-EDTA). Infiltrated leaves were placed into the buffer containing the spin trap reagents and illuminated for 30 min with white light (100 μmol quanta m^−2^s^−1^). At the end of the illumination time, the leaves were removed and the EPR signal of the solution was monitored. EPR spectra were recorded at room temperature in a standard quartz flat cell using an ESP-300 X-band spectrometer (Bruker). The following parameters were used: microwave frequency 9.73 GHz, modulation frequency 100 kHz, modulation amplitude: 1 G.

### Analysis of aldehydes, including reactive electrophile species and β-cyclocitral, with LC-MS/MS

Aldehydes were measured according to Roach et al., (2017). Briefly, leaves frozen in liquid nitrogen were ground for 1 min inside 2 mL reaction tubes using 2 × 5 mm quartz beads in 1 ml of pre-cooled (−20 °C) acetonitrile, containing 0.5 μM 2-ethylhexanal (as internal standard) and 0.05 % (w:v) of butylated hydroxytoluene. The resulting extract was centrifuged for 10 min, 4°C, at 26,000 *g*, and the supernatant was split with 600 μL used for the analysis of aldehydes and 200 μL for the measurement of chlorophyll. Aldehydes were derivatized with 2,4-Dinitrophenylhydrazine (2,4-DNPH) in the presence of formic acid and diluted 50:50 with LC-MS-grade H_2_O before injection of 3 μL sample. Separation was carried out using a reversed-phase column (NUCLEODUR C18 Pyramid, EC 50/2, 50×2 mm, 1.8 μm, Macherey-Nagel, Düren, Germany), using an ekspert ultraLC 100 UHPLC system coupled to a QTRAP 4500 mass spectrometer (AB SCIEX, Framingham, MA, USA). Peak areas of selected ions were normalised to 2-ethylhexanal and chlorophyll content of the sample. Chlorophyll was quantified by absorbance in 80% acetone using the extinction coefficients of Porra et al. (1989).

### Carotenoid measurement

Mature leaves were frozen in liquid nitrogen, ground and lyophilized. Powder (2 mg dry weight) was dissolved in 500 μl acetone, 0,01% ammoniac, then centrifugation at 12000 *g* 5 min, supernatants were collect and pellet washed twice in acetone. Solvent was evaporated and the precipitates were resuspended in 100 μl acetone. Liquid chromatography-UV-absorption (HPLC-UV) analysis was performed using a HPLC column (Uptisphere Strategy C18-HQ 250 × 3 mm, 3 μm granulometry, Interchim, Montluçon, France) with a HPLC/UV chain (Shimadzu, Tokyo, Japan) including two pumps (LC-20AD), a sample manager (SIL-20AC HT), a column oven (CTO-20A) and an UV diode array detector (UVSPD-M20A) with a flow of 0.5 mL min^−1^. The mobile phase A was composed of 10% water and 89.5% acetonitrile with 0.5% acetic acid and the mobile phase B was ethyl acetate with 0.5% acetic acid. The elution gradient was as follows: initially 10% B, 1 min 10% B; then a linear increase up to 95% B for 24 min. At the end, the column was returned to initial conditions for a 15 min equilibration. Absorbance was monitored at 450 nm. Signals were calculated by normalization of peak areas to chlorophyll *a*.

### Statistical analysis

Graphs and statistical tests were produced using the R software. For all boxplots, the various elements are defined as follows: upper whisker = largest observation less than or equal to upper hinge + 1.5 × interquartile range (IQR); lower whisker = smallest observation greater than or equal to lower hinge – 1.5 × IQR; center line=median, 50% quantile; upper hinge = 75% quantile; lower hinge = 25% quantile; outliers are data beyond the end of the whiskers and represented by a dot; each dot in the box represents single data points. Symbols represent the p-value of Student test ns (non significant); *: p-value ≤ 0.05; **: p-value ≤ 0.01; ***: p-value ≤ 0.001; ****: p-value ≤ 0.0001.

## Results

To investigate the drought stress tolerance of state transition mutants, we subjected wild type, *nsi1, nsi2*, and *stn7* to drought stress by withholding water for six days. The wild type leaves wilted while the state transition mutants were visibly much less affected (Fig. 1A). For the following experiments water was withhold for four days to induce moderate drought stress. Under this condition, photosynthetic electron transport is not considered the primary damage site (Kaiser 1987; Cornic and Fresneau 2002). Upon moderate drought stress, the rosettes of the mutants showed a much higher water content (Fig. 1B). Furthermore, stomata were significantly more closed in wild type upon moderate drought stress, while there was no change in stomata opening in the mutants (Fig. 1C).

**Fig. 1.**
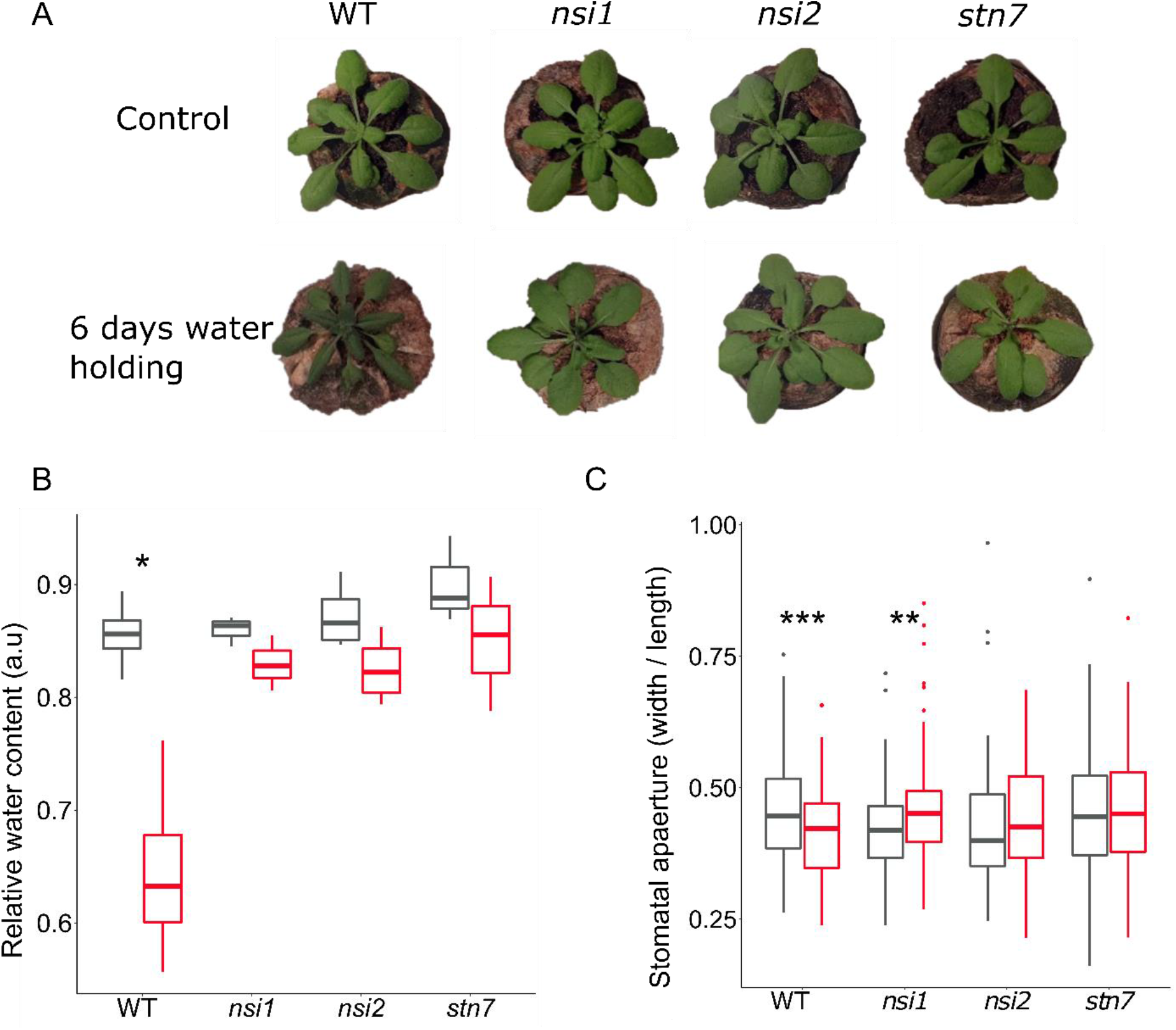
State transition mutants are more resistant to drought. **A:** 4-week-old *A. thaliana* wild type and *nsi1, nsi2, stn7* plants well-watered and after 6 days without watering. **B:** Relative water content of total shoot in control conditions (grey) and after 4 days without watering (red). **C:** Stomata aperture. Ratio width/length of stomata aperture in control conditions (black) and after 4 days without watering (red), counting more than 83 stomata on at least 3 different leaves in each condition. Three replicates from four different independently grown sets of plants, n=12, mean and SD are given. Stars indicate significant differences according to Student’s t-test (* p<0.05, ** p<0.01, *** p<0.001).

Since the mutants are affected in their ability to adapt the antenna size dependent on the light quality, intensity, and stress conditions, we investigated the effect of moderate drought on their photosynthesis using chlorophyll fluorescence. No significant differences in pigment composition were observed between the genotypes (SI Table 1). Analysis of chlorophyll fluorescence parameters showed that upon drought the effective quantum yield of photosystem II (YII) was affected in wild type while there were no significant changes in the mutants (Fig. 2A). For the statistical analysis, values measured at a light intensity of 100 μmol quanta m^−2^s^−1^ were chosen, an intensity close to the growth light intensity. When the light intensity exceeds the optimal intensity for assimilation, energy dissipation at the level of PSII sets in. According to Kramer et al. (2004), two types of quantum yields for energy dissipation can be defined, Y(NPQ) and Y(NO) which represent regulated and non-regulated energy dissipation, respectively. Dissipation of excess energy expressed as the yield of non-photochemical quenching, Y(NPQ), increased in the wild type upon drought while it remained constant in the mutants (Fig. 2B). Relevant here is that state transitions have only a minor contribution to chlorophyll fluorescence measurements of Y(NPQ), thus lack of change of Y(NPQ) in the mutants reflects lack of stress. This was the case of the regulated part of NPQ while the non-regulated quenching, Y(NO), decreased in the wild type upon stress (Fig. 2C). When the genotypes were compared in control conditions, Y(NO) was significantly higher in the mutants, showing that state transitions influence chlorophyll fluorescence at growth light intensity. The larger amount of LHCII at PSII has the consequence that reaction centers of PSII were more closed in the mutants as indicated by significantly lower qL values in the control conditions, i.e., the primary quinone acceptor Q_A_ accumulated more in its reduced form in the mutants. In the case of qL, there was a significant decrease in the wild type upon drought, showing that more reaction centers were closed, while no differences were observed for the mutants (Fig. 2H).

**Fig. 2.**
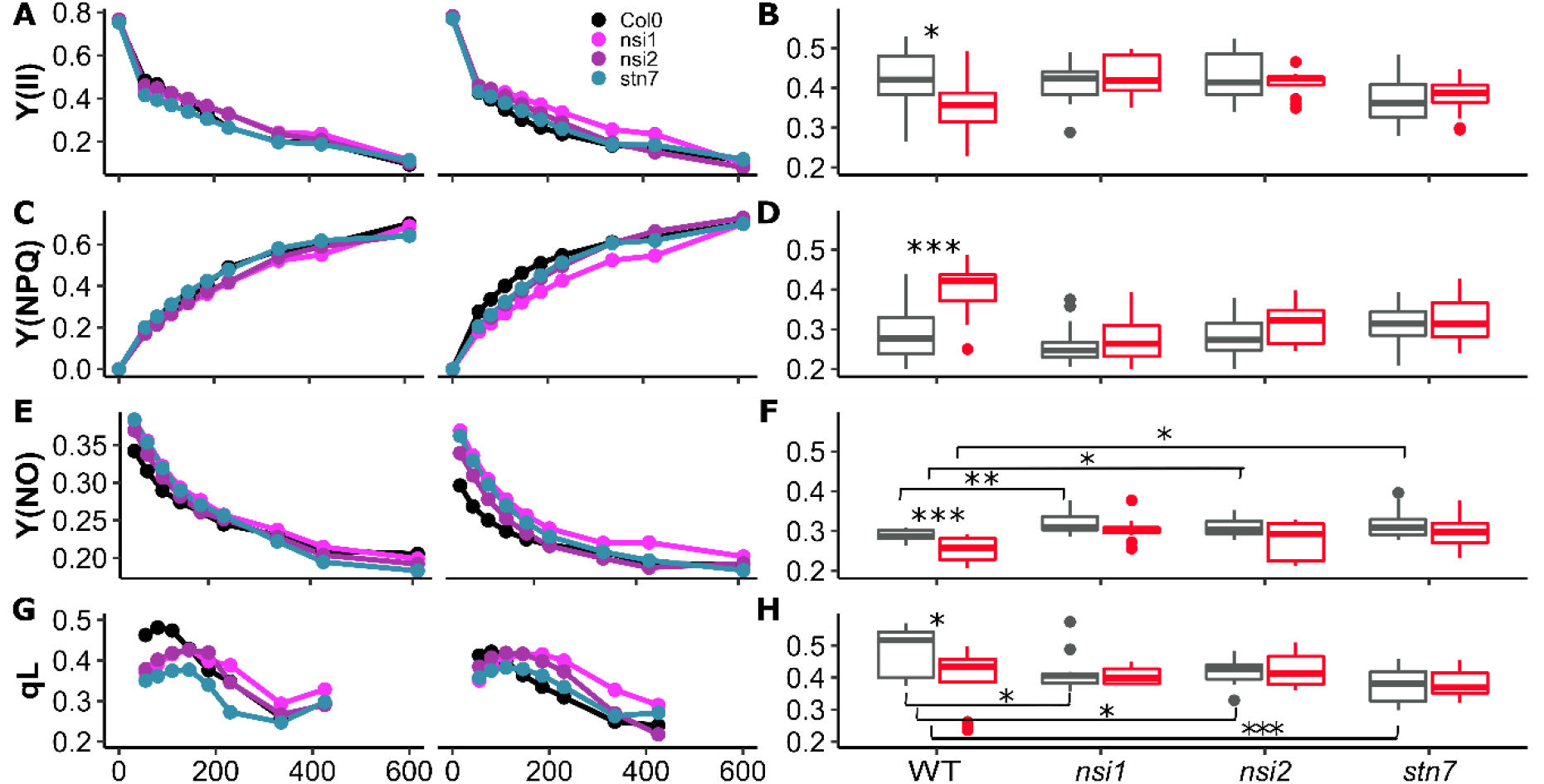
State transition mutants show no changes in the light dependency of chlorophyll fluorescence parameters upon moderate drought stress. Left: light response curves of fluorescence parameters in control conditions or after 4 days without watering (stress). **A, B:** effective quantum yield PSII, Y(II), **C, D:** controlled non-photochemical quenching, Y(NPQ), **E, F:** Non-regulated non-photochemical quenching, Y(NO), and **G, H:** fraction of open PSII centers, qL. The left panel shows variations of the fluorescence parameters as a function of the light intensity; the right panel shows Box plots of the fluorescence parameters for wild type and *nsi1, nsi2, stn7* at growth light intensity (100 μmol m^− 2^s^− 1^). Each light intensity was applied for 3 min before giving a saturating light flash. Three replicates from four different independently grown sets of plants, n=12, mean and SD are given. Stars indicate significant differences according to Student’s t-test (* p<0.05, ** p<0.01, *** p<0.001).

The results of the chlorophyll fluorescence parameters are in line with a lack of state transition in the mutants, however, they do not explain their improved drought tolerance. Since it has been described previously that N-acetylation of protein is important for germination and root development (Linster et al., 2015), we investigated the root growth of the seedlings. As shown in Fig. 3A, B, the main root of the wild type was thinner and shorter compared with the mutants. In addition, lateral root development was strongly enhanced in the mutants. The difference between the lengths of the main root in wild type and mutants was even higher when plants were grown under osmotic stress conditions (200 mM mannitol) (Fig. 3C). However, when the photosynthetic electron transfer was blocked by DCMU, an inhibitor that blocks electron transport in PSII by binding to the Q_B_-site of the D1 protein, or by DBMIB, an inhibitor of the cytochrome *b*_6_*f* complex that binds to the Qo-binding site, differences between the genotypes were no longer observed. In the presence of DBMIB that leads to a reduced PQ pool, growth of the main root was even stimulated in all genotypes compared to control conditions, while growth was retarded in the presence of DCMU that leads to an oxidized PQ pool. These results indicate that the reduction state of the PQ pool controls root growth. When seedlings were grown in the dark, no significant differences in hypocotyl or root length were observed (Fig. 3F, G), demonstrating that photosynthetic electron transport reactions are responsible for the difference between the wild type and the state transition mutants.

**Fig. 3:**
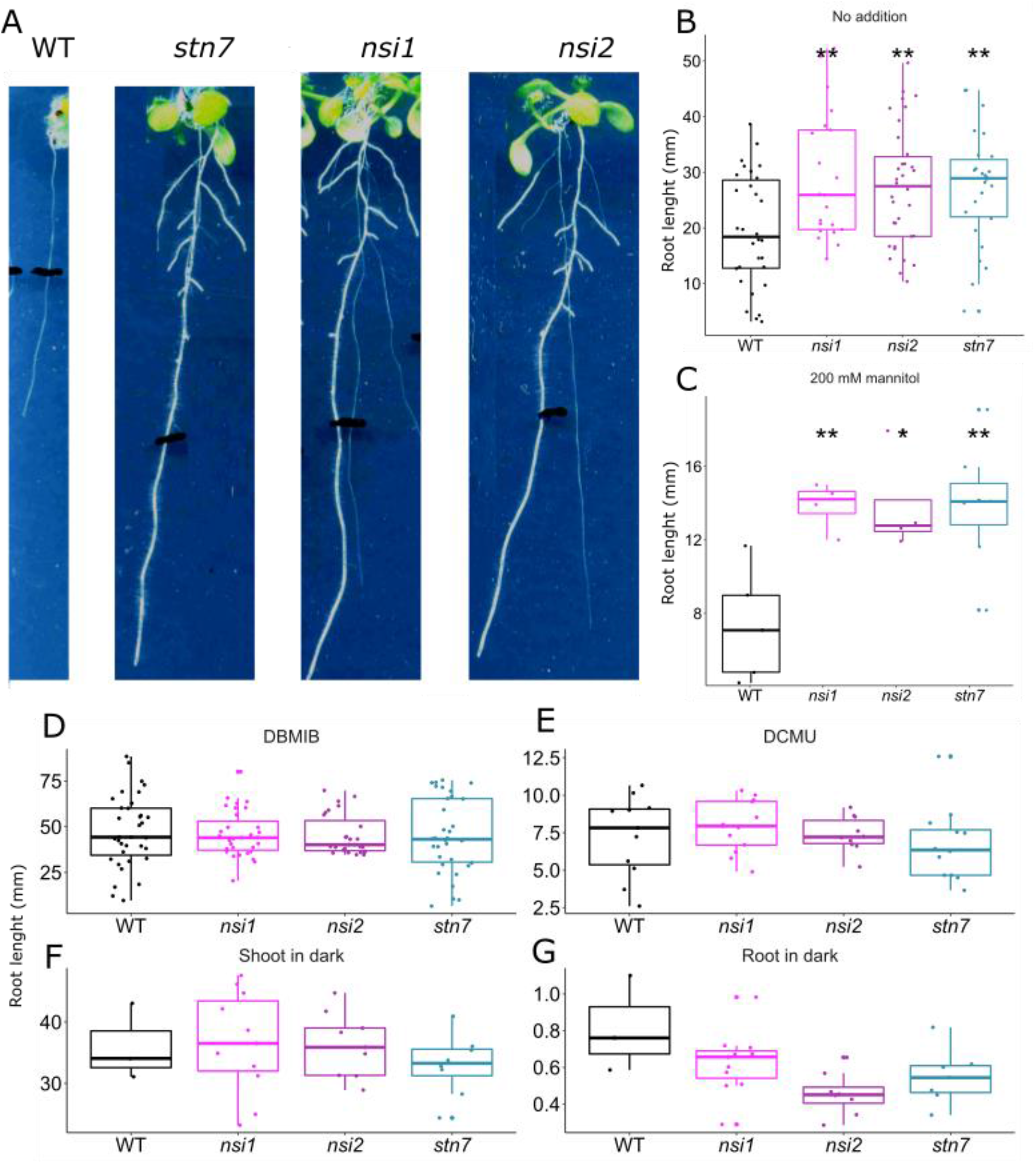
State transitions mutants show stimulation of growth root. **A:** Photos of seedings of WT, *nsi1, nsi2*, and *stn7* after 10 days of growth, black lines have been drawn at the end of the roots after 7 days. **B:** Length of the primary root of 7 days old in control conditions, n ≥ 21. **C:** Length of the primary root of 7 days old in the presence of 200 mM mannitol. **D** and **E:** Length of the primary root of 7 days old in the presence of 30 μM DBMIB, n ≥ 10, and 2 μM DCMU respectively, n ≥ 10. **F and G:** Length of the shoot and root of 7 days old in dark, n ≥ 17 and 14 respectively. Mean and SD are given. Stars indicate significant differences according to Student’s t-test (* p<0.05, ** p<0.01).

In excess light conditions, more light is absorbed by the antenna than can be used for the reduction of NADP^+^, and the PQ pool becomes more reduced. This is the case in the state transition mutants as seen by the lower qL compared to the wild type (Fig. 2 G, H). When Q_A_ is reduced, charge recombination reactions within PSII do occur between Q_A_^−^and the oxidized primary donor P680^+^. This reaction leads to the generation of ^1^O_2_, a highly oxidizing reactive oxygen species. For visualizing ^1^O_2_ generation in leaves, we first used the fluorophore Singlet Oxygen Sensor Green, but, using this approach, we were not able to quantify differences between the four genotypes. Instead, we used an EPR spin probing assay with TEMPD as a probe to detect ^1^O_2_ (Hideg et al., 2008). The assay cannot be performed with leaves or intact chloroplasts, and we used isolated thylakoid membranes instead. In thylakoids from wild-type plants, the L-LHCII is partly localized at PSI as is the case in leaves, while in the mutants all LHCII is localized at PSII (Fig. 4A). As expected, when the PQ pool is more oxidized, less ^1^O_2_ was generated in wild type compared with the mutants (Fig. 4B, C).

**Fig. 4:**
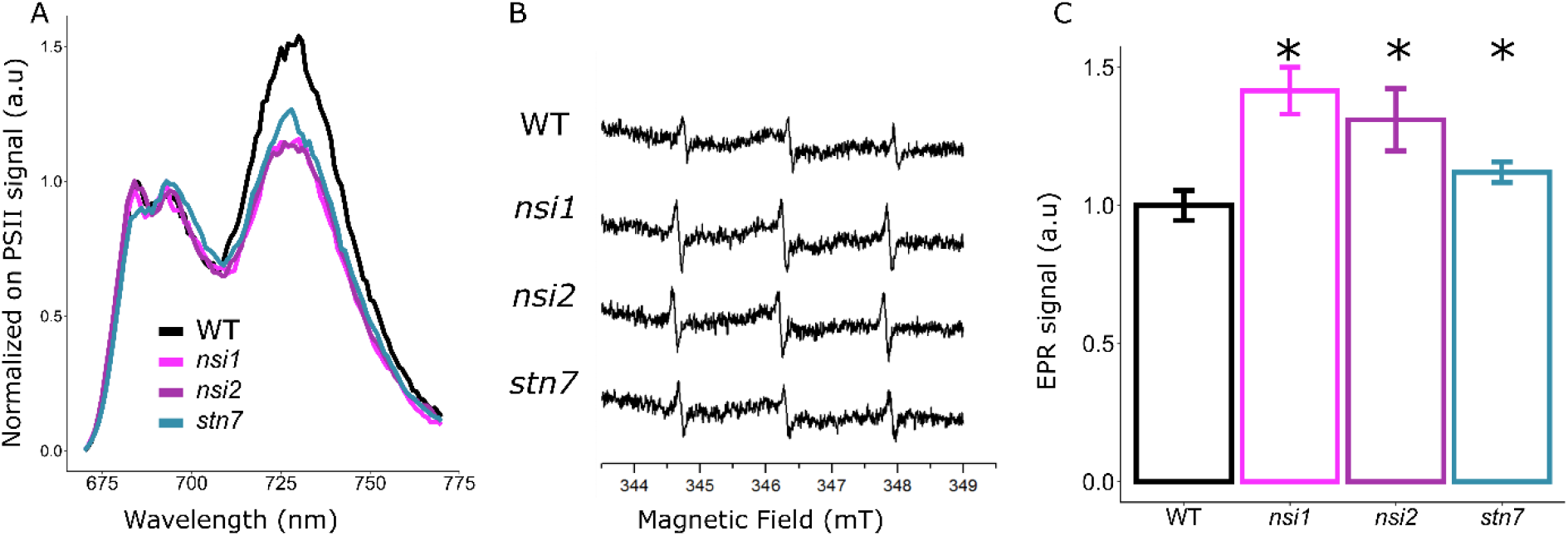
Thylakoid membranes of state transition mutants generate more singlet oxygen. **A:** 77 K chlorophyll fluorescence emission spectra of isolated thylakoid membranes. Signal normalised to PSII emission at 695 nm. Representative spectra are shown (n=3). **B:** Detection of singlet oxygen by EPR using the spin probe TEMPD-HCl. Typical spectra are shown. **C:** Quantification of the EPR signals (n=4, 2 thylakoid preparations from set of plants grown at different times). Mean and SD are given. Stars indicate significant differences according to Student’s t-test ** p<0.01).

To investigate whether there is a general increase in ROS generation in state transition mutants, we measured O_2_^• –^/H_2_O_2_-derived hydroxyl radicals by a spin-trapping assay with 4-POBN/EtOH as a trap (Mubarakshina et al., 2010). Furthermore, the activity of a few antioxidant enzymes was measured. The amounts of O_2_^•–^/H_2_O_2_ and the activities of the antioxidant enzymes superoxide dismutase, peroxidase, and catalase were the same in all genotypes (SI Fig. S1), showing that there was not a general increase in the level of oxidative stress in the mutants, but instead, a specific rise of ^1^O_2_ levels. In the PSII reaction center, ^1^O_2_ can react with β-carotene, leading to oxidation products like β-cyclocitral and β-ionone. In addition, ^1^O_2_ can react with lipids giving rise to additional reactive electrophile species (RES). Fig. 5 shows the most abundant RES detected in leaves from the wild type and the state transition mutants. In control conditions, there is no significant difference between the genotypes (Fig. 5A), while upon moderate drought stress the wild type generated significantly more β-cyclocitral, β-ionone, acrolein, hydroxyhexanal, and butyraldehyde (Fig. 5B). This is typical for a plant submitted to stress conditions. This shows that the wild-type plants are highly stressed, as already visible by the eye, while the state transition mutants cope with moderate drought stress without showing stress symptoms (Fig. 1A).

**Fig. 5:**
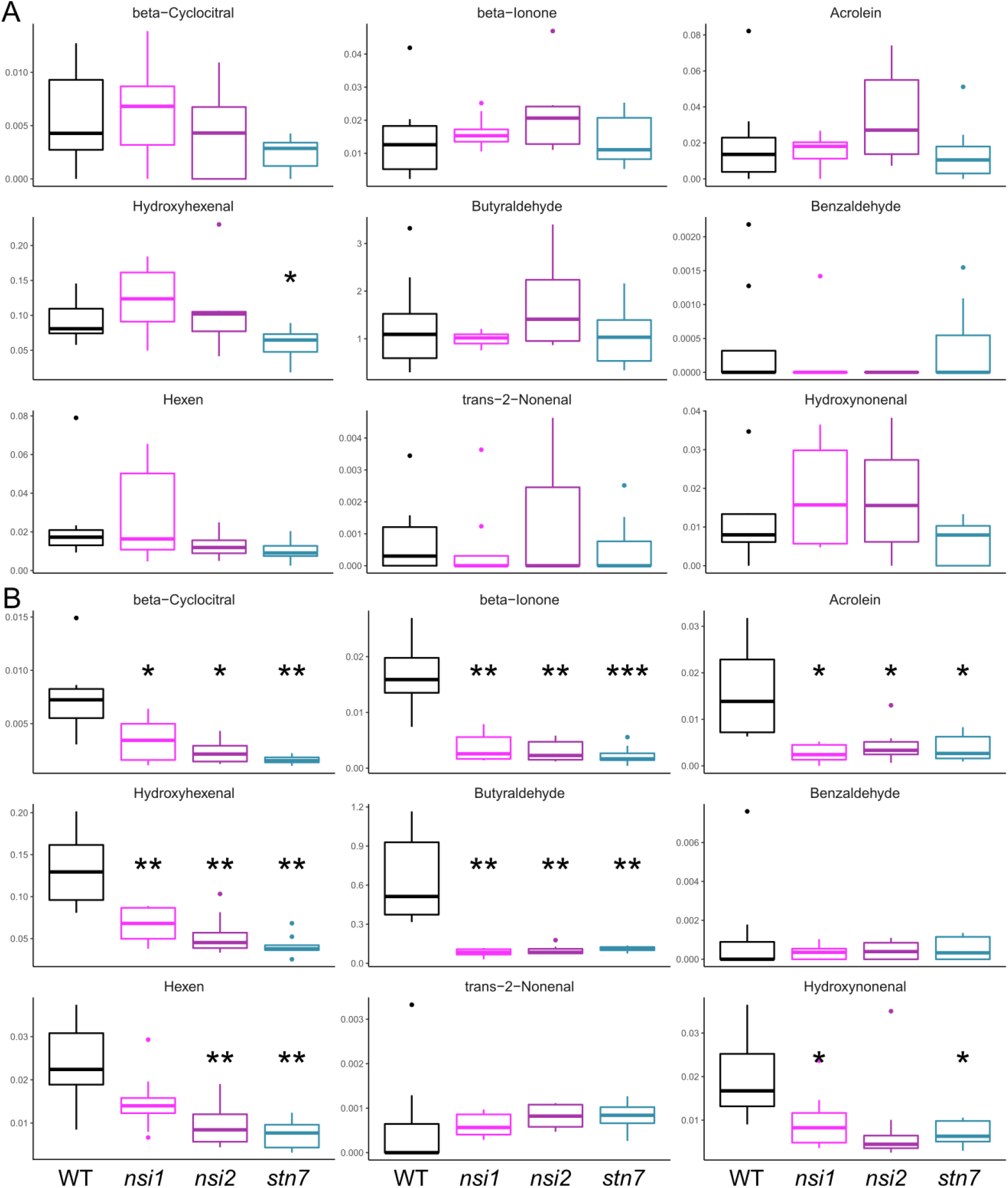
Generation of reactive electrophile species is stimulated in wild type upon moderate drought stress. **A:** Amounts of RES in leaves of 4-week-old WT, *nsi1, nsi2*, and *stn7* in control conditions. Shown are ion peak areas relative to the internal standard and normalized to the chlorophyll content of the extract. **B:** Same as A after 4 days without watering (n=8, independently grown sets of plants). Mean and SD are given. Stars indicate significant differences according to Student’s t-test (* p<0.05, ** p<0.01, *** p<0.001).

## Discussion

State transitions are controlled by the reduction state of the PQ pool via the activation of the STN7 kinase (Depège et al., 2003; Bellafiore et al., 2005). The results reported here raise the question of how the reduction state of the PQ pool affects root development. According to the data shown in Fig. 3, the reduction state of the plastoquinone pool is crucial for primary root growth and lateral root development. In state transition mutants, the PQ pool is more reduced than in the wild type (Fig.2H). Moreover, drought typically stimulates root growth, which also associates with a more reduced PQ pool as seen in the wild type (Fig. 2H). In all genotypes, when the PQ pool is reduced in the presence of DBMIB, root growth is stimulated (Fig. 3D), whereas shorter roots are observed when the PQ pool is oxidized in the presence of DCMU (Fig. 3E). Both inhibit photosynthetic electron transport, thereby showing that it is indeed the redox state of the PQ pool that exerts control of root growth and not differences in photosynthesis and sugar production. The inhibitory effect of DCMU on primary and lateral root growth has been observed previously and has been interpreted as photosynthesis promoting lateral root emergence partly through auxin biosynthesis (Duan et al., 2021). However, in the study by Duan and coworkers, the inhibitor DBMIB has not been used. Since both DCMU and DBMIB inhibit photosynthesis, we rule out differences in auxin levels being involved in the early root architecture phenotype we observe here.

As shown in Fig. 4, state transition mutants generate slightly more ^1^O_2_ than the wild type. ^1^O_2_ may be the key to signaling events that promote root growth. Addition of DCMU inhibited root growth (Fig. 3E), speaking at a first glance against the hypothesis that ^1^O_2_ is responsible for the stimulation of root growth. However, although the addition of DCMU favors charge recombination in PSII, the yield of ^1^O_2_ is low thanks to the modification of the midpoint potential of the redox couple Q_A_/Q_A_^−^favoring charge recombination between P680^+^ and Q_A_^−^via a direct route that does not yield ^3^Chl (Krieger-Liszkay and Rutherford, 1998; Fufezan et al., 2002). The redox state of the PQ pool is known to be important for retrograde signaling (Pfalz et al., 2012; Dietz et al., 2016). In retrograde signaling, a signal travels from the chloroplast to the nucleus within the same cell. Here, a signal is required that travels over long distances. ^1^O_2_ may produce signals such as apocarotenoids that are able to initiate signaling events far from the site of their production. β-carotene oxidation products such as the volatile β-cyclocitral and its direct water-soluble oxidation product, β-cyclocitric acid, are known to regulate nuclear gene expression through several signaling pathways (Ramel et al., 2012; Shumbe et al., 2017; D’Alessandro et al., 2019). However, we were not able to detect an increase in β-cyclocitral or other RES in leaves of state transition mutants compared to wild type in plants under control conditions. This may be because the difference in ^1^O_2_ production was, although significant, small. Even smaller differences are expected in the subsequently generated β-carotene oxidation products. We may technically not be able to detect such small differences in RES. Differences in levels of β-cyclocitral or other RES may also depend on the developmental stage of the plants. The RES measurements were performed with mature leaves while the root phenotype was observed in young seedlings. Besides ^1^O_2_ and RES, the redox state of the PQ pool may activate other signaling pathways that involve miRNAs (Bertolotti et al., 2021) or changes in hormone levels like abscisic acid and strigolactones favoring root growth (Gomez–Roldan et al., 2008; Ruyter-Spira et al., 2011).

In conclusion, alterations in state transition appear to be a promising trait for improving plant growth and thereby crop productivity under harsh environmental conditions. The signaling pathway has to be elucidated in future work. It should be explored whether other photosynthetic mutants which exhibit an imbalance between PSII and PSI activity are also more drought resistant. Mutants affected not only in the antenna systems but also in the activity of the photosystems seem to be promising avenue for exploring drought tolerance. Especially mutants with slight defects in the turnover of the cytochrome *b*6*f* complex or in photosystem I may be of interest since their PQ pool is in a more reduced state. Mutants lacking small subunits, for example PsaI (Schöttler et al., 2017), or expressing only one isoform of certain subunits, for example PsaE1 or PsaE2 (Hald et al., 2008; Krieger-Liszkay et al., 2020), which do not show a strong phenotype in control conditions seem to be ideal candidates to test their resistance to drought stress.

## Acknowledgements

We would like to thank Sandrine Cot (I2BC) for technical assistance and Paula Mulo (University of Turku, Finland) for sending us the seeds of the state transition mutants. This work was supported by the Labex Saclay Plant Sciences-SPS (ANR-17-EUR-0007), the platform of Biophysics of the I2BC supported by the French Infrastructure for Integrated Structural Biology (FRISBI; grant number ANR-10-INSB-05), this work has benefited from the support of IJPB’s Plant Observatory technological platforms and this work benefited from the French state aid managed by the ANR under the “Investissements d’avenir” programme with the reference ANR-16-CONV-0003 (CLand). L.L. is supported by a CLand and SPS PhD fellowship.

## Author Contribution

L.L., and A.K-L. designed the project. L.L., T.R., F.P., and A.K-L. performed the experiments and analysed the data. F.M. participated in discussions. L.L. and A.K-L. wrote the initial version of the manuscript that was read and revised by all authors.

### Data availability

Data will be made available on demand.

