## Supplemental Table for "Increased drought resistance in state transition mutants is linked to modified plastoquinone pool redox state"

**Supplemental Table S1.**

**Determination of pigments in leaves of plants in control conditions or after 4 days without watering (stress)**

| **Control** | | | | |
|---|---|---|---|---|
| | WT | nsi1 | nsi2 | stn7 |
| **Chl B** | 3.963±0.162 | 3.933±0.191 | 4.182±0.188 | 3.963±0.208 |
| **β-carotene** | 0.797±0.075 | 0.767±0.083 | 0.798±0.076 | 0.809±0.094 |
| **Lutein** | 1.295±0.134 | 1.234±0.139 | 1.316±0.127 | 1.301±0.164 |
| **Violaxanthin** | 0.035±0.005 | 0.038±0.005 | 0.04±0.004 | 0.034±0.006 |
| **Zeaxanthine** | 0.02±0.007 | 0.02±0.006 | 0.018±0.005 | 0.017±0.006 |
| **Antheraxanthin** | 0.014±0.004 | 0.061±0.038 | 0.047±0.026 | 0.021±0.009 |
| **Neoxanthin** | 0.079±0.016 | 0.061±0.016 | 0.068±0.017 | 0.07±0.02 |
| **Stress** | | | | |
| **Chl B** | 0.068±0.025 | 0.082±0.031 | 0.103±0.029 | 0.075±0.018 |
| **β-carotene** | 3.988±0.223 | 4.154±0.302 | 4.379±0.266 | 4.294±0.263 |
| **Lutein** | 0.029±0.009 | 0.017±0.006 | 0.034±0.008 | 0.024±0.008 |
| **Violaxanthin** | 0.125±0.052 | 0.188±0.072 | 0.183±0.084 | 0.098±0.038 |
| **Zeaxanthine** | 0.067±0.073 | 0.082±0.074 | 0.084±0.079 | 0.083±0.101 |
| **Antheraxanthin** | 0.032±0.005 | 0.031±0.006 | 0.044±0.005 | 0.041±0.008 |
| **Neoxanthin** | 1.462±0.187 | 1.48±0.217 | 1.501±0.199 | 1.661±0.235 |

Quantification by LC-MS measurement on leaves of 4-week-old WT, *nsi1*, *nsi2*, and *stn7* in control conditions and after 4 days of water holding for stress. Signal is area of pics for each species normalized by area of Chl A pic. At least 2 matures leaves of 2 different plants were collected for the 8 technical replicates for each genotypes and conditions.

**Supplemental Figure 1**

**Activity of antioxidant enzymes in crude protein extracts from leaves in control and moderate stress conditions**

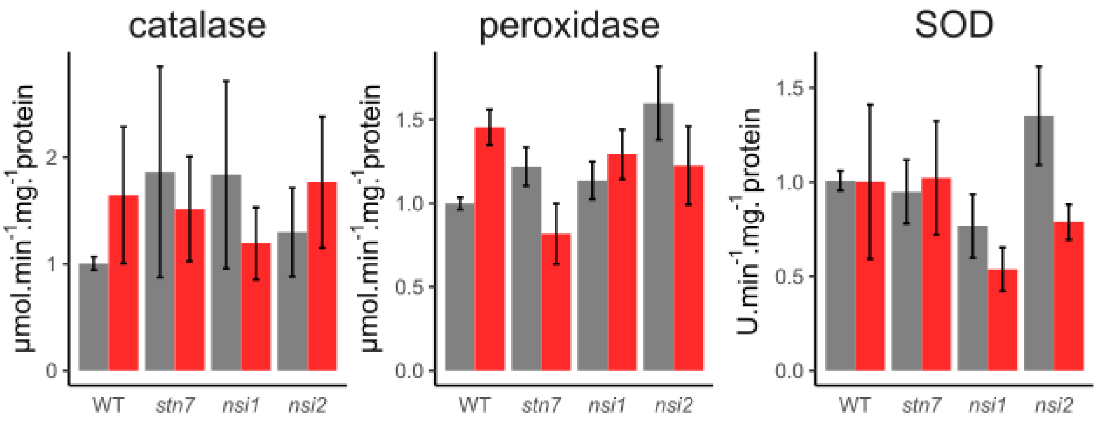


Activities were normalised to the activity of WT plants in control condition. A crude protein extracts were prepared by grinding leaves in liquid nitrogen before homogenization in a buffer containing 0.3 M sorbitol, 50 mM Kcl, 5 mM $MgCl_2$, 20 mM HEPES pH 7.2. The supernant after centrifugation was used for the measurements. Protein content was determined using Amioblack. Enzyme activities were determined according to Molins et al. 2013. Grey bars: well-watered; red: plants were not watered for 4 days. N≥9, 3 different extractions from plants grown at different times. Mean ± SD are shown. Activities WT control catalase 2.85 µmol.min-$^{1}$.mg.protein-$^{1}$; peroxidase 133 µmol.min-$^{1}$.mg.protein-$^{1}$.µmol.min and 632 U.min-$^{1}$.mg.protein-$^{1}$ for superoxide dismutase.
